# Life origin and evolution may be influenced by the dynamical Coupling between Earth’s magnetic field strength and atmospheric oxygen level

**DOI:** 10.1101/2025.08.13.670079

**Authors:** Maodi Liang, Qinghua Cui

## Abstract

A recent study has revealed that Earth’s magnetic field strength and atmospheric oxygen level exhibit strongly correlated, linearly increasing trends (M-O relationship) over the past 540 million years. This discovery sheds new lights on the long-standing debate regarding whether a strong magnetic field is an essential prerequisite for origin and the evolution of complex life. Here, we demonstrate that the M-O relationship is not static but highly dynamic across geological timescales using dynamic correlation and variable parameter regression analysis. Intriguingly, periods of weakened M-O coupling coincide precisely with the “Big Five” mass extinctions, while the strongest M-O correlation aligns with the Cambrian explosion. Further analysis indicates that shifts in M-O linkage are driven primarily by changes in the magnetic field’s trend rather than oxygen fluctuations, with the “Big Five” extinctions closely matching inflection points in Earth’s magnetic field strength curve. Additionally, human gene age analysis unveils that *de novo* gene birth rate correlates strongly with mass explosion/extinction and M-O relationship strength, enabling the prediction of such events predating 540 million years ago (Mya) and potentially in the future. Our findings suggest that the stability of Earth’s magnetic field trend—or its coupling strength with oxygen—rather than absolute magnetic strength or oxygen level, plays a pivotal role in shaping the rise and fall of life.

## INTRODUCTION

Earth is the only planet known in the Universe confirmed to host life(*1*) with a documented biological history extending back at least 3.77 billion years(*2*). What enables Earth to sustain life, and how does it maintain long-term habitability? This stands as one of the most fundamental questions in modern science, with profound implications for understanding life’s origin and evolution**—**as well as guiding the search for extraterrestrial life(*1*). In addition to fundamental requirements such as liquid water, appropriate temperature ranges, and stable gravity, scientists have proposed several other critical factors for Earth’s long-term habitability. Among these, Earth’s strong intrinsic magnetic field has been identified as a potentially essential element(*3*), supported by temporal correlations between paleomagnetic records and the emergence of life. However, current research findings remain inconsistent and often contradictory, highlighting the need for more comprehensive investigations into the geomagnetic field’s influence on planetary habitability(*1*).

It is well established that variations in the strength and direction of Earth’s geomagnetic field throughout the planet’s history can be reconstructed by analyzing rocks formed by ancient volcanic eruptions(*4*). Similarly, indirect methods allow us to estimate atmospheric oxygen level over the past ∼540 million years(*5*). Recently, Kuang et al. combined two key datasets— the virtual geomagnetic axial dipole moment (VGADM) reconstructed from paleomagnetic records and atmospheric oxygen level inferred from geochemical proxies— to uncover a striking correlation between Earth’s magnetic field strength and atmospheric oxygen level(*1*). These findings demonstrate that over the past 540 million years, both Earth’s magnetic field strength and atmospheric oxygen level exhibit a strongly correlated, linearly increasing trends (referred to as the M-O relationship). While the underlying mechanism driving this puzzling M-O relationship remains unclear, this discovery offers new insights into the long-standing debate on whether a strong magnetic field is an essential prerequisite for the origin and evolution of complex life(*6*). Furthermore, this mysterious link may provide a fresh perspective for understanding Earth’s long-term habitability.

On the other hand, it is well known that there have been five big mass extinction events in Earth’s history (called the ‘Big Five’)(*7*), which was first identified by Raup and Sepkoski through investigating the marine fossil record in 1982(*8*). Over the past four decades, numerous theories, causes, and mechanisms have been proposed to explain these events(*9*), for example bolide impacts, large igneous province eruption, volcanic eruptions(*10*), tectono-oceanic changes, and astronomical triggers (e.g. solar flares, gamma bursts and supernova explosions), however, none fully explain all events with definitive evidence(*11*). The causes of mass extinctions remain one of the greatest mysteries(*11*), demanding fresh perspectives and critical thinking(*11*).

In this study, by analyzing the M-O relationship at higher temporal resolution using dynamic correlation and variable parameter regression, we reveal that the M-O correlation exhibits significant variability, ranging from strongly positive to negative or even statistically insignificant. This demonstrates that the M-O relationship is not static but dynamically variable across geological timescales. Notably, periods of weakened M-O coupling coincide precisely with the “Big Five” mass extinctions, whereas the strongest M-O correlation aligns with the Cambrian explosion. Further analysis suggests that shifts in M-O linkage are primarily driven by changes in the magnetic field’s trend rather than oxygen fluctuations, with the “Big Five” extinctions closely corresponding to inflection points in Earth’s magnetic field strength curve. Additionally, human gene age analysis reveals that *de novo* gene birth rates correlate strongly with mass explosion/extinction events and M-O link strength, enabling the prediction of such events predating 540 million years ago (Mya) and potentially in the future. Based on these findings, we infer two previously unrecognized mass explosions (∼1488 and 987 Mya) and three mass extinctions (∼1934, 1292, and 810 Mya) predating 540 Mya. Our findings suggest that the stability of Earth’s magnetic field trend—or the strength of its coupling with oxygen— rather than absolute magnetic strength or oxygen level, plays a pivotal role in driving the prosperity and decline of species.

## RESULTS

### The dynamic relationship between Earth’s magnetism and atmospheric oxygen level

As revealed by Kuang et al., Earth’s magnetic field strength and atmospheric oxygen level exhibit a strongly correlated, linearly increasing trends over time (M-time: Rho = 0.60, p-value = 6.10e-55, **Figure 1A**; O-time: Rho = 0.69, p-value = 9.86e-77, **Figure 1B**; M-O correlation: Rho = 0.79, p-value = 1.74e-115). To investigate the M-O relationship at a higher temporal resolution, here we conducted a sliding-window Spearman’s correlation analysis (see Methods). Our results reveal that the M-O relationship is not static but instead highly dynamic across geological timescales (Supplemental File S1, **Figure 1C**). For instance, from 540 and 449 Mya, the M-O correlation was strongly positive, but it declined to non-significance after 449 mya, turned strongly negative by 439 Mya, and reverted to non-significance again after 382 Mya (**Figure 1C**). Additionally, the M-O relationship at 1-million-year resolution revealed by VPR analysis further demonstrates it is not static but highly dynamic (Supplemental File S2, **Figure 2A**). These findings suggest that although the global trend in M-O correlation has been positive over the Phanerozoic Con, the relationships exhibit significant variability at finer temporal scales.

**Figure 1.**
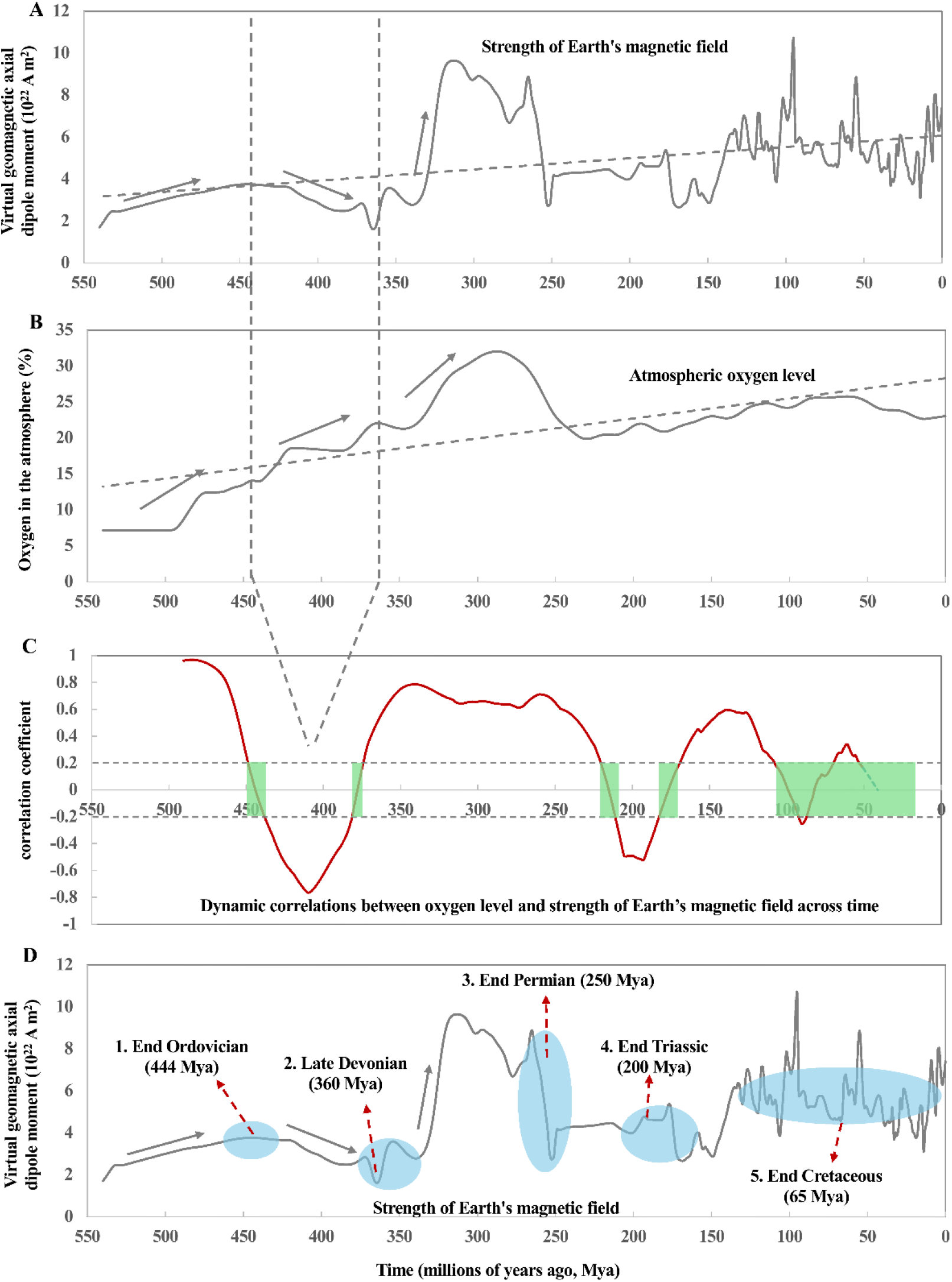
Temporal dynamics of Earth’s magnetic field strength, atmospheric oxygen levels, and their relationship (M-O link) over the past 540 million years. (**A**) Earth’s magnetic field strength and (**B**) atmospheric oxygen levels both exhibit long-term increasing trends. (**C**) Dynamic correlation analysis reveals the M-O relationship is not static but shows significant temporal variability, with correlation coefficients (Rho) ranging from strongly positive to negative. The gray-shaded region between Rho values of 0.2 and -0.2 indicates statistically non-significant correlations (bounded by dotted horizontal lines). The period between vertical dotted lines in A and B illustrates a characteristic example of dynamic M-O coupling. (**D**) Analysis demonstrates that variations in M-O correlation strength are driven primarily by inflection points in Earth’s magnetic field strength trend rather than changes in atmospheric oxygen levels. Notably, periods of weakened M-O correlation (green-shaded regions in **C**) and magnetic field trend inflection points (light blue-shaded regions in **D**) show precise temporal correspondence with known mass extinction events.

**Figure 2.**
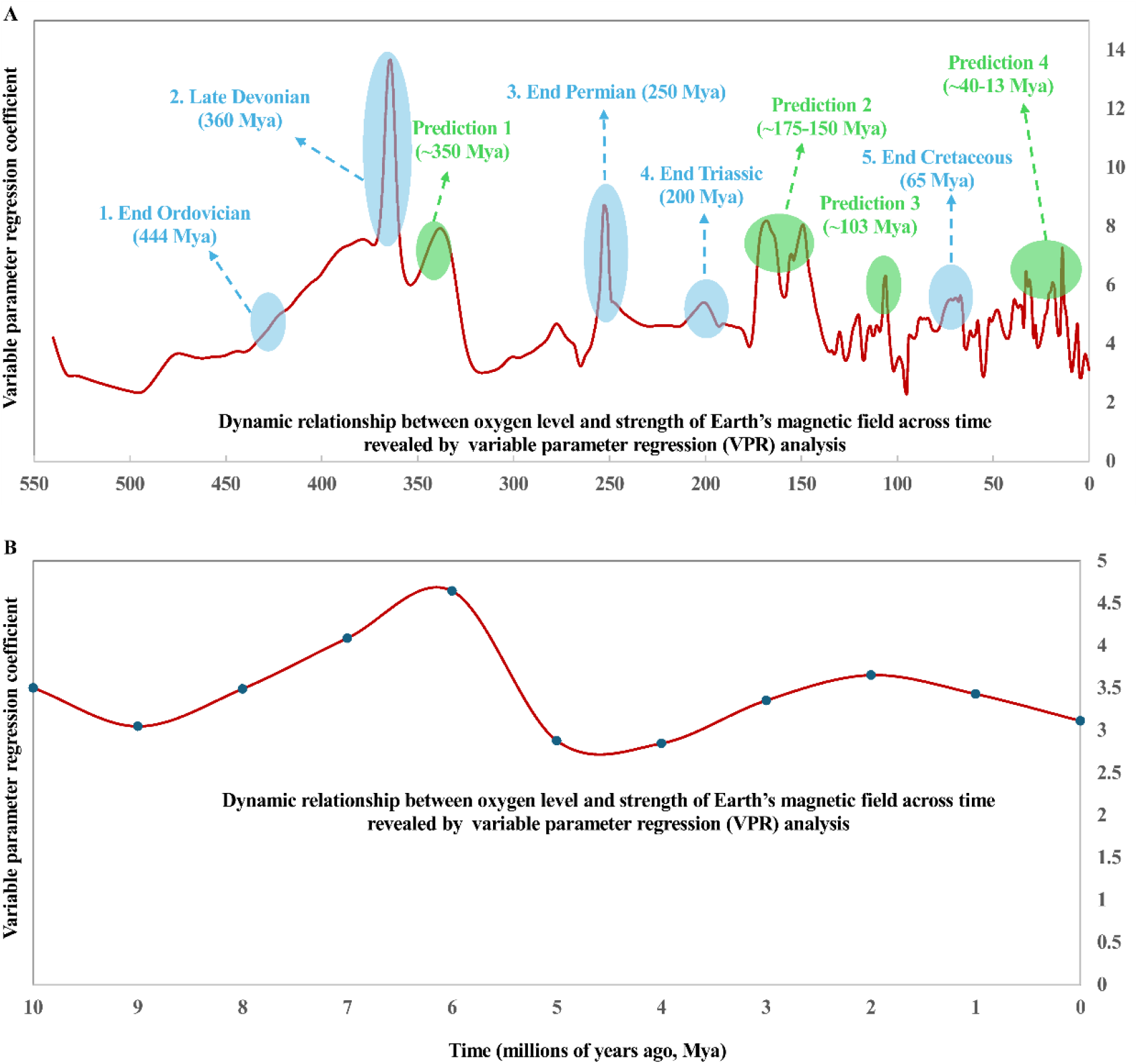
Variable parameter regression (VPR) analysis of the dynamic M-O relationship. (**A**) High-resolution (1-million-year timescale) reconstruction of the M-O relationship over the past 540 million years. Sharp increases in the VPR coefficient (β > 5.0) show precise temporal correspondence with both the canonical “Big Five” mass extinction events (light blue-shaded regions) and four newly identified extinction events (green-shaded regions). (**B**) Detailed view of the M-O relationship dynamics during the most recent 10 million years, demonstrating the continued variability of this coupling at finer timescales.

Given these dynamic patterns, it is crucial to explore how positive, negative, and non-significant M-O correlations may have influenced the origin and evolution of life on Earth throughout geological history. Our dynamic correlation analysis revealed that the M-O relationship weakened—becoming statistically non-significant—during the periods coinciding with the ‘Big Five’ mass extinctions (green-shaded regions in **Figure 1C**). This finding was further corroborated by VPR analysis, which revealed sharp increases in the VPR coefficient (*β* > 5.0) precisely aligning with these five mass extinction events: the End Ordovician (444 Mya), Late Devonian (360 Mya), End Permian (250 Mya), End Triassic (200 Mya), and End Cretaceous (65 Mya) extinctions (light blue-shaded regions in **Figure 2A**).

A critical question then arises: which factor—magnetic field strength or atmospheric oxygen level—primarily drives the variability in the M-O relationship? As clearly illustrated in **Figure 1**, the dynamics of M-O correlation are predominantly governed by changes in magnetic field strength rather than atmospheric oxygen level. For instance, while both magnetic field strength (**Figure 1A**) and atmospheric oxygen level (**Figure 1B**) showed increasing trends from 540 to 449 Mya, their trajectories diverged markedly thereafter. After 449 Mya—a key inflection point in the magnetic field strength curve (**Figure 1D**) —the magnetic field strength began to decline while the atmospheric oxygen level continued to rise (region between the dotted vertical lines in **Figure 1A-B**). These inflection points in the magnetic field strength trend directly correspond to transitions in the M-O relationship strength (**Figure 1C**), alternative between positive, non-significant, and negative correlations. Notably, these inflection points exhibit remarkable temporal coincidence with the ‘Big Five’ extinction events quite well (light blue-shaded regions in **Figure 1D**).

### Birth rate of human *de novo* genes corresponds to the M-O relationship strength

Current evidence suggests life originated approximately 3.8 billion years ago(*12*). However, since the fossil record of macroscopic life only becomes rich from ∼540 Mya, onward, our understanding of mass extinction events is currently limited to his more recent period. This raises several fundamental questions: Did mass extinctions occur prior to 540 Mya (here termed “B540 mass extinctions”)? How can we identify evidence for such ancient mass extinction event? Are we currently experiencing a sixth mass extinction? Might there be additional mass extinctions beyond the established “Big Five”? More importantly, can we predict future trends in mass explosions and extinctions? These questions remain both critically important and persistently challenging. Our discovery of the dynamic M-O relationship and geomagnetic field strength trend changes offers a novel framework to address these issues.

To investigate mass explosions and extinctions across a deep time, here we propose a new hypothesis, periods of mass explosions or extinctions may correlate with high or low number of *de novo* gene births, respectively. To test this, we analyzed the **n**umber of human ***d****e novo* **g**enes of specific **a**ges (NDGA; Supplementary File S3). Strikingly, NDGA shows remarkable correspondence with known mass explosion and extinction events (**Figure 3A**). Notably, each of the “Big Five” extinction events coincides with reduced *de novo* gene birth rates, while the Cambrian explosion period exhibits the highest number of *de novo* gene origins (**Figure 3A**). This finding enables us to predict pre-540 Mya mass explosion and extinction events through NDGA analysis. For instance, only 12 genes originated around 1934 Mya, compared to 1,211 genes at 1488 Mya. It is clear that there are three valleys and three peaks in the *de novo* gene number curve before 540 Mya (**Figure 3B**). The three peaks (∼1488, ∼987, and ∼645 Mya) likely represent mass explosions, with the most recent (∼645 Mya) potentially corresponding to the Cambrian explosion (541-530 Mya). Given the temporal resolution limits of gene age estimation (the next datapoint after 645 Mya is 544 Mya, with 1,586 *de novo* genes), we propose that ∼645-544 Mya represents an extended “super explosion” period encompassing the fossil-record-defined Cambrian explosion(*13*). This temporal offset suggests fossil evidence may lag behind the actual initiation of major evolutionary events. Consequently, we identify two additional predicted explosion events (∼1488 and ∼987 Mya) and three mass extinction events (∼1934, 1292, and 810 Mya) before 540 Mya.

**Figure 3.**
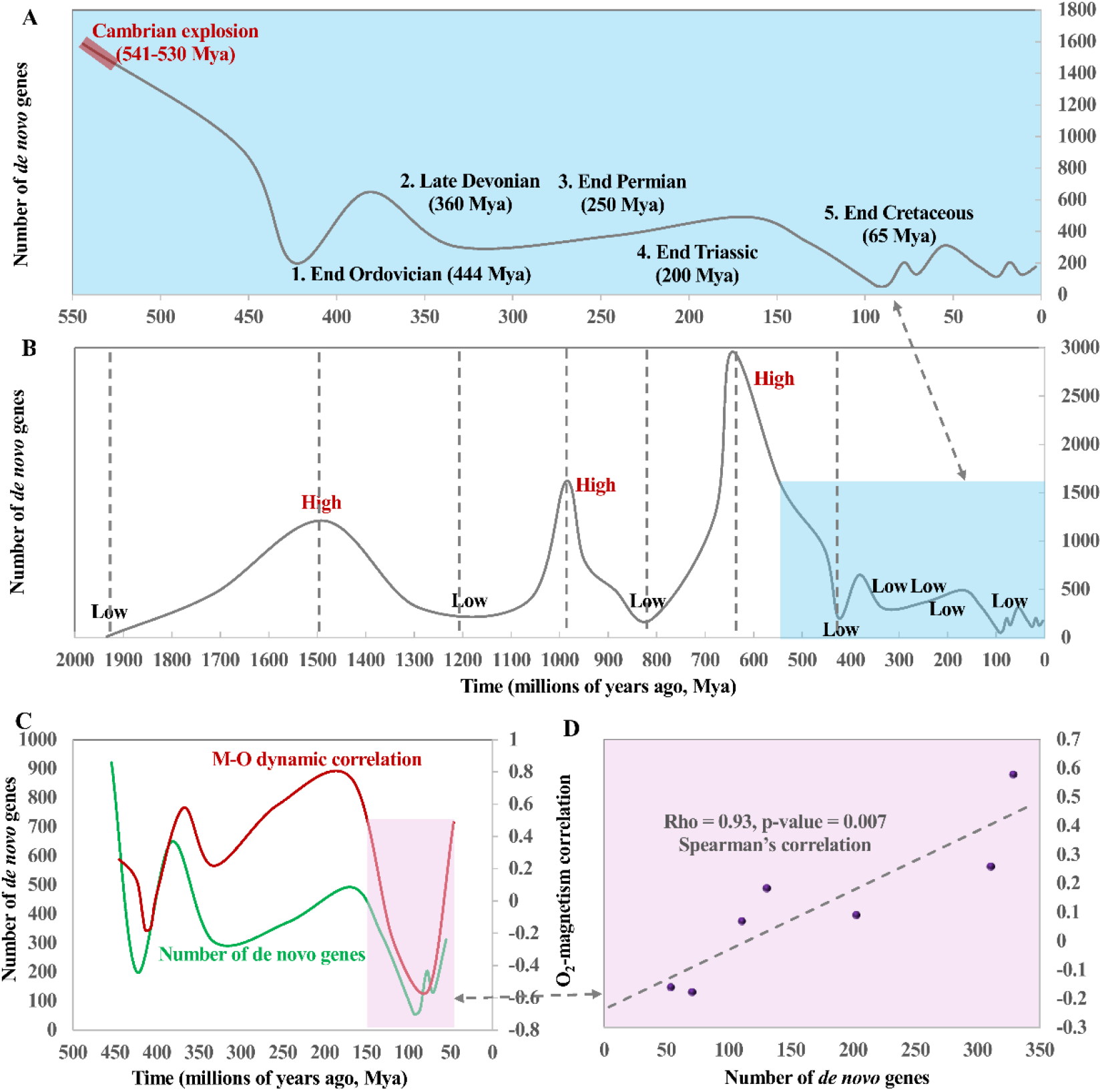
Correlation between human *de novo* gene birth rates, mass evolutionary events, and the M-O relationship. (A) Temporal distribution of human *de novo* gene origins over the past 540 million years shows striking correspondence with both the “Big Five” mass extinction events and the Cambrian explosion. (B) Extension of this analysis to pre-540 million years reveals three predicted mass extinction events (valleys) and two mass explosion events (peaks) based on *de novo* gene birth rates. (C-D) Significant positive correlation between *de novo* gene birth rates and the strength of the M-O relationship, demonstrating their coupled dynamics through geological time.

The strong correspondence between NDGA and mass events suggests a potential linkage with M-O dynamics. Indeed, NDGA and M-O correlation trends show striking similarity over the past 540 Mya (**Figure 3C**), particularly during the last 200 Mya where they exhibit a strong positive correlation (Rho=0.93, p-value = 0.007, **Figure 3D**).

Another question is that whether there are additional mass extinction events besides the established “Big Five” over the past 540 million years. Our high-resolution (1-million-year time scale) VPR analysis reveals four previously unrecognized potential mass extinction events (∼350, ∼175-150, ∼103 Mya, and ∼40-13 Mya; green-shaded regions in **Figure 2A**) alongside the “Big Five”. While the ∼350 Mya event (Prediction 1) may correspond to the Late Devonian extinction in lower-resolution fossil records, the distinct temporal separation (∼10 My) at our resolution suggests these represent separate events. This demonstrates the enhanced sensitivity of our approach compared to traditional methods.

Presently (0 Mya), the VPR coefficient remains below the critical threshold (*β* > 5.0) and shows a declining trend (**Figure 2B**), suggesting that we are not currently in—nor immediately facing—a sixth mass extinction at million-year timescales. However, considering longer (100 Mya) timescales, Earth’s magnetic field strength exhibited intense oscillations accompanied by persistently weak M-O correlations (**Figure 1C-D**), suggesting we may be experiencing an extended mass extinction event spanning the last 100 million years.

## DISCUSSION

Understanding whether and how Earth’s magnetic field serves as an essential prerequisite for the origin and evolution of complex life represents a fundamental scientific question. Equally critical is elucidating the mechanisms underlying mass evolutionary explosions and extinctions. Our study reveals that the relationship between Earth’s magnetic field strength and atmospheric oxygen levels (M-O relationship) is dynamic rather than static, as demonstrated through dynamic correlation and variable parameter regression analyses. This M-O correlation exhibits substantial temporal variability, ranging from strongly positive to negative or statistically insignificant values.

Notably, we establish that three key indicators—the dynamic M-O relationship, trends in Earth’s magnetic field strength, and human *de novo* gene birth rates—collectively predict mass extinction and explosion events with remarkable precision, including the canonical “Big Five” mass extinctions and the Cambrian explosion. Based on these relationships, we identify two previously unrecognized mass explosions (∼1488 and 987 Mya) and three ancient mass extinctions (∼1934, 1292, and 810 Mya). Crucially, our findings demonstrate that the stability of Earth’s magnetic field trend and its coupling strength with atmospheric oxygen—rather than absolute values of either parameter—plays a determining role in species proliferation and decline. These insights provide transformative perspectives on two longstanding questions: the role of geomagnetic fields in life’s history, and the fundamental drivers of mass extinction events.

Our findings enable several important predictive applications. Reconstruction of paleomagnetic and atmospheric oxygen data beyond 540 Mya could permit prediction of ancient mass events through our dynamic analytical framework. Such predictions could then be cross-validated with *de novo* gene birth rate analyses. However, we note current limitations in gene age estimation methods, particularly for ancient genes, highlighting the need for improved bioinformatics approaches to achieve higher temporal resolution.

Given that it is the trend change of Earth’s magnetic field strength but not is absolute value play a vital role in the prosperity and decline of species, it is emergently important to search for the mechanisms causing the change of Earth’s magnetic field strength. The birth rate analysis of human *de novo* genes revealed that the mass explosions and extinctions clearly fluctuate in a periodic mode (as pointed by the dotted grey lines in **Figure 3B**). As of 400 Mya, the birth rate curve shows 4 valleys and peaks (1934, **1488**, 1292, **987**,810,**645**, and 424 Mya, the peaks are highlighted in bold). Interestingly, the average interval between a valley and its neighbor peak is 252 My, which is strikingly close to a galactic year, which is estimated to be 225-250 My, defined as the time it takes for our entire Solar System to complete one orbit around the center of Milky Way. This temporal correspondence suggests a potential astrophysical influence on Earth’s magnetic field dynamics, where specific orbital positions may trigger geomagnetic trend changes that subsequently drive biotic turnover. Based on this speculation, it is not difficult to predict that there was a mass explosion event around the period of 172 Mya after the mass extinction event of 424 Mya (424-252 = 172). Indeed, there is a peak (168 Mya, 492 *de novo* genes; **Figure 3B**) of de novo gene birth around 172 Mya. Extending this pattern, we predict an impending mass extinction event approximately 84 million years in the future (168 - 252 = -84). These findings open new avenues for investigating influences on Earth’s biosphere dynamics across geological timescales.

## MATERIALS AND METHODS

### Data used in this study

We obtained the datasets of Earth’s magnetic field strength and atmospheric oxygen level over the past 540 million years from Kuang et al.’ study(*1*). Data regarding the “Big Five” mass extinctions events were obtained from Our World in Data (https://ourworldindata.org/mass-extinctions)(*14*). The age of human genes, that is, the origin time of as *de novo* genes, was retrieved from the GenOrigin database(*15*). To keep precision, we removed 939 genes with uncertain *de novo* birth age (all annotated as “>4290”, Supplemental File S4). Finally, 16,385 human genes with information of *de novo* birth age were used in this study (Supplemental File S5).

### Data analysis

To examine the M-O relationship at higher temporal resolution, we conducted dynamic Spearman’s correlation analysis using a sliding window approach. Employing a 100-million-year window that advanced in 1-million-year increments, we initially analyzed the period from 540 to 441 million years ago (Mya). The window was then systematically shifted (539-440 Mya) and the correlation repeated, with this process continuing iteratively until reaching the final window (99-0 Mya).

Additionally, we investigated the M-O relationship at a finer temporal resolution (1-million-year) using variable parameter regression (VPR) analysis, which we previously proposed to quantify dynamic gene connectivity based on temporal gene expression profiles(*16*). The VPR framework is defined as follows:

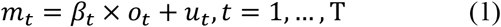

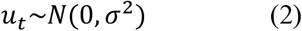

where *m*_*t*_ and *o*_*t*_ represent Earth’s magnetic field strength and atmospheric oxygen level at time *t*, respectively; *β*_*t*_ is the time-varying regression coefficient, reflecting the M-O coupling strength at time *t*; *u*_*t*_ denotes Gaussian distribution with zero mean and σ standard deviation. Then, the dynamic variation of *β* over time is given by:

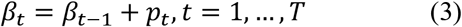

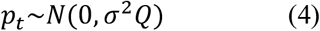

where *σ*^2^*Q* is the stationary covariance matrix of the innovation *p*_*t*_. Using this framework, we estimated the dynamic M-O relationship at single time-point resolution (1-million-year scale), capturing its fine-scale variability across geological time.

All statistical analyses were performed using R software (version 4.4.3).

## Acknowledgements

We thank our colleagues for valuable discussion.

## Author contributions

Writing—original draft: Q.C. Conceptualization: Q.C. Investigation: M.L., Q.C. Supervision: Q.C.

